# Mutational Analysis and Deep Learning Classification of Uterine and Cervical Cancers

**DOI:** 10.1101/2022.10.12.511895

**Authors:** Paul Gomez

## Abstract

We analyzed tumor mutations of 7 uterine and 2 cervical cancers with the goal of developing a Deep Learning (DL) software tool that can automatically classify tumors based on their somatic mutations. The data were obtained from the AACR Genie Project [1], that has a collection of more than 120,000 tumor samples for more than 750 cancer types. We performed a thorough analysis of the mutational data of tumors of the uterus and uterine cervix, selecting tumors with 3 or more mutations and cancer types with more than 15 cases. For each cancer type we then selected the top 12 most mutated genes among their neoplasms. In the introduction section we summarize our analysis of these nine diseases and in the deep learning section we present a convolutional neural network (CNN) [2] that yields an overall classification accuracy of 94.3% and 89.2% on the train and test datasets, respectively. We hope that this tool can be added to the existing arsenal of histological and immunohistochemical techniques in cases when a precise diagnosis cannot be clearly determined. Each cancer type has a unique somatic mutational profile that can be used to disambiguate two candidate malignancies with similar histologic features.

## INTRODUCTION

### Uterine Cancers

This year, 2022, approximately 66,000 patients in the United States are estimated to be diagnosed with uterine or endometrial cancer [3]. The number of uterine cancer patients worldwide was 417,000 in 2020. Uterine cancer is the fourth most common cancer for women in the United States. It is estimated that in 2022, approximately 12,550 patients will die of uterine cancer [3], making it the sixth most deadly cancer among women in the United States.

More than 90% of uterine cancers occur in the endometrium. Endometrial cancers are classified as Type I (endometrioid subtype) or Type II (non-endometrioid subtype) [4][5]. The differences between the two groups lie on precursor type, unopposed estrogen presence, menopausal status, myometrial invasion, histologic subtypes, and genetic mutations [6][7].

Type I neoplasms of the uterus are low grade tumors that start with a precursor lesion called atypical hyperplasia (AH) [8][9] that develops in premenopausal patients in the presence of unopposed estrogen, that is, in the absence of progesterone. Endometrial hyperplasia is the proliferation of glands of irregular size and shape with a high gland-to-stroma ratio [10][11]. Endometrial hyperplasia can be cytological atypical or non-atypical [12][13]. The presence or absence of nuclear atypia is the main feature to determine if a carcinoma is of Type I. AH lesions show none or low myometrial invasion and thus, they are confined to the endometrium. The most common carcinoma of this type is Endometrial Carcinoma (UCEC) [14][15] (See Table 1 and Figure 1).

**Table 1.** Uterine and Cervical Cancers

| Organ | Code | Disease Name | Uterine Type | Cancer Type | Tissue/Histologic subtype |
| --- | --- | --- | --- | --- | --- |
| Cervix | CESC | Cervical Squamous Cell Carcinoma |  | Carcinoma | Squamous cell |
| Cervix | ECAD | Endocervical Adenocarcinoma |  | Adenocarcinoma | Glandular epithelium |
| Uterus | UCCC | Uterine Clear Cell Carcinoma | Type II | Carcinoma | Clear Cell |
| Uterus | UCEC | Endometrial Carcinoma | Type I | Carcinoma | Endometrium |
| Uterus | UCS | Uterine Carcinosarcoma/Uterine Malignant Mixed Mullerian Tumor | Type II | Sarcoma | Myometrium, müllerian |
| Uterus | UEC | Uterine Endometrioid Carcinoma | Type I | Carcinoma | Endometrioid |
| Uterus | ULMS | Uterine Leiomyosarcoma |  | Sarcoma | Myometrial |
| Uterus | UMEC | Uterine Mixed Endometrial Carcinoma | Type II | Carcinoma | Mixed subtypes |
| Uterus | USC | Uterine Serous Carcinoma/Uterine Papillary Serous Carcinoma | Type II | Carcinoma | Serous |

**Figure 1.**
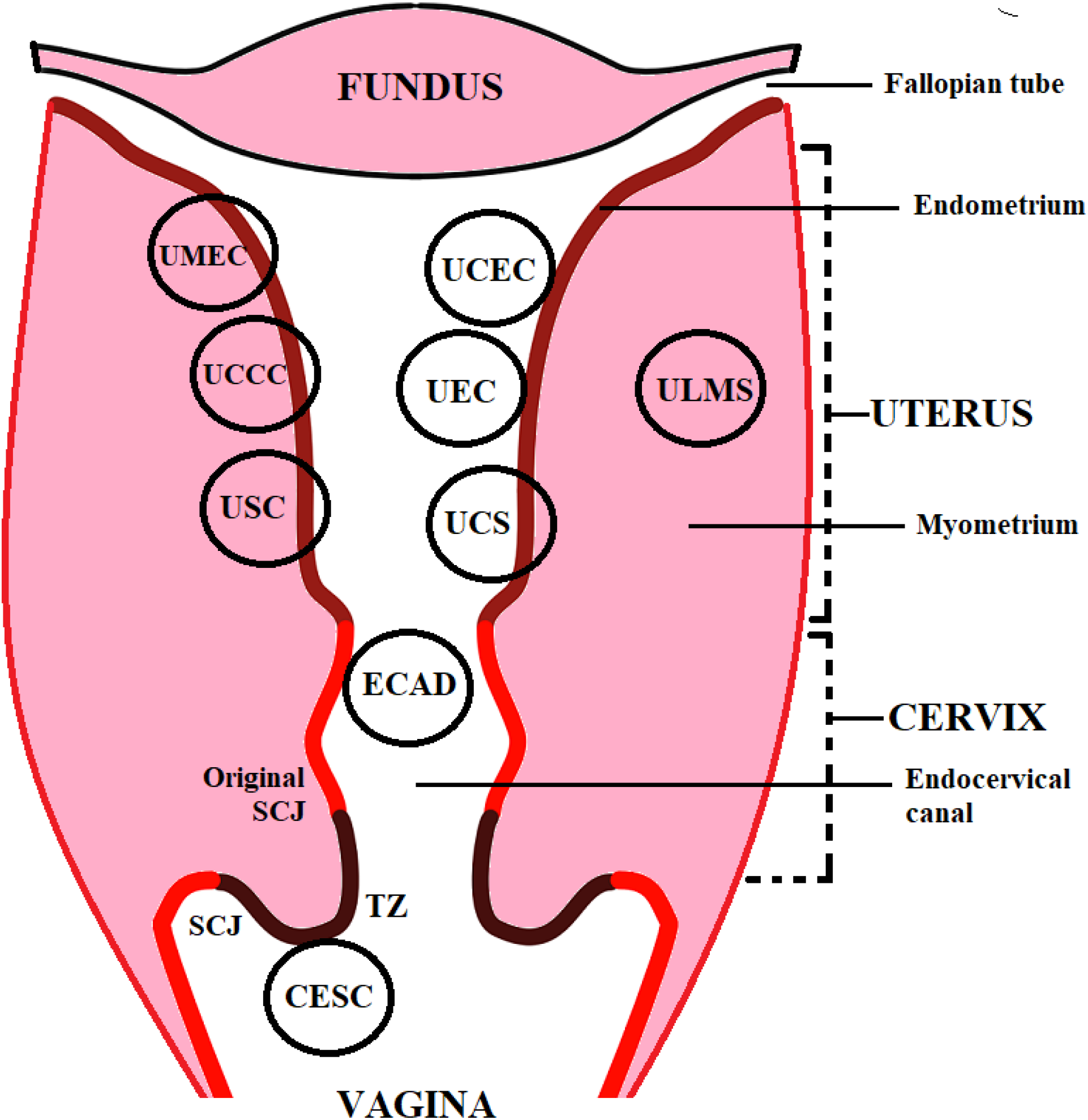
Diagram showing the relative locations of neoplasms of the uterus and uterine cervix.

At the molecular level, mutations of gene PTEN have been identified as an initial driver of tumorigenesis in all hyperplasias and endometrioid neoplasms [16][17][18][19]. PTEN is a tumor suppressor gene involved in a signal transduction path that regulates cell growth and apoptosis [19][20]. On Table 2, it can be seen that all uterine cancers except Uterine Leiomyosarcoma (ULMS) have PTEN mutated. ULMS is a sarcoma that does not fall in any of the Type I or Type II categories. ULMS is a rare cancer of the uterus [22][23] that was included in this study due to its unique mutational pattern having 3 unique mutated genes. These ULMS unique genes, namely, DAXX, ERBB4, and KDR, are not present on the mutational profiles of the other cancers on Table 2.

**Table 2.**
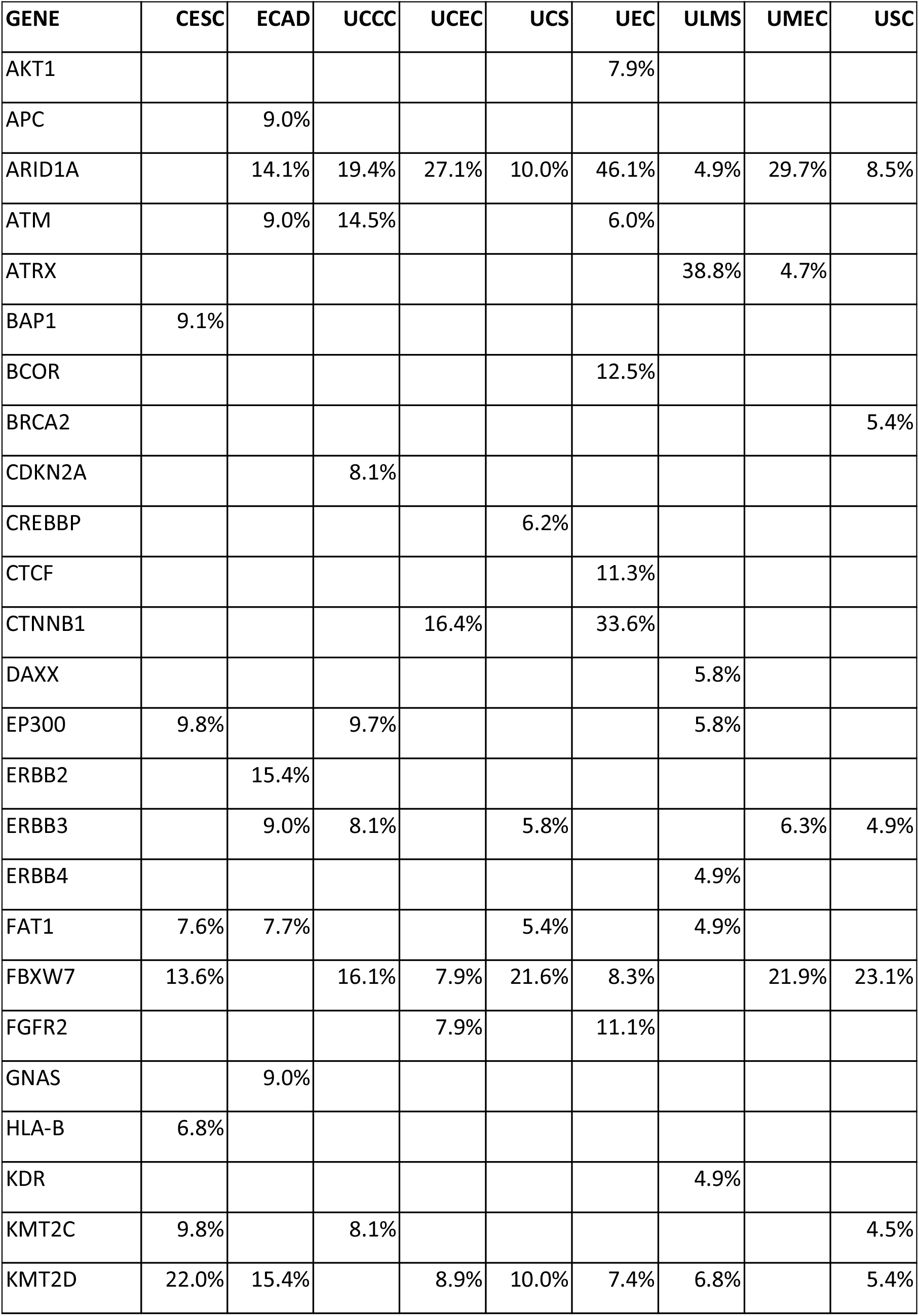
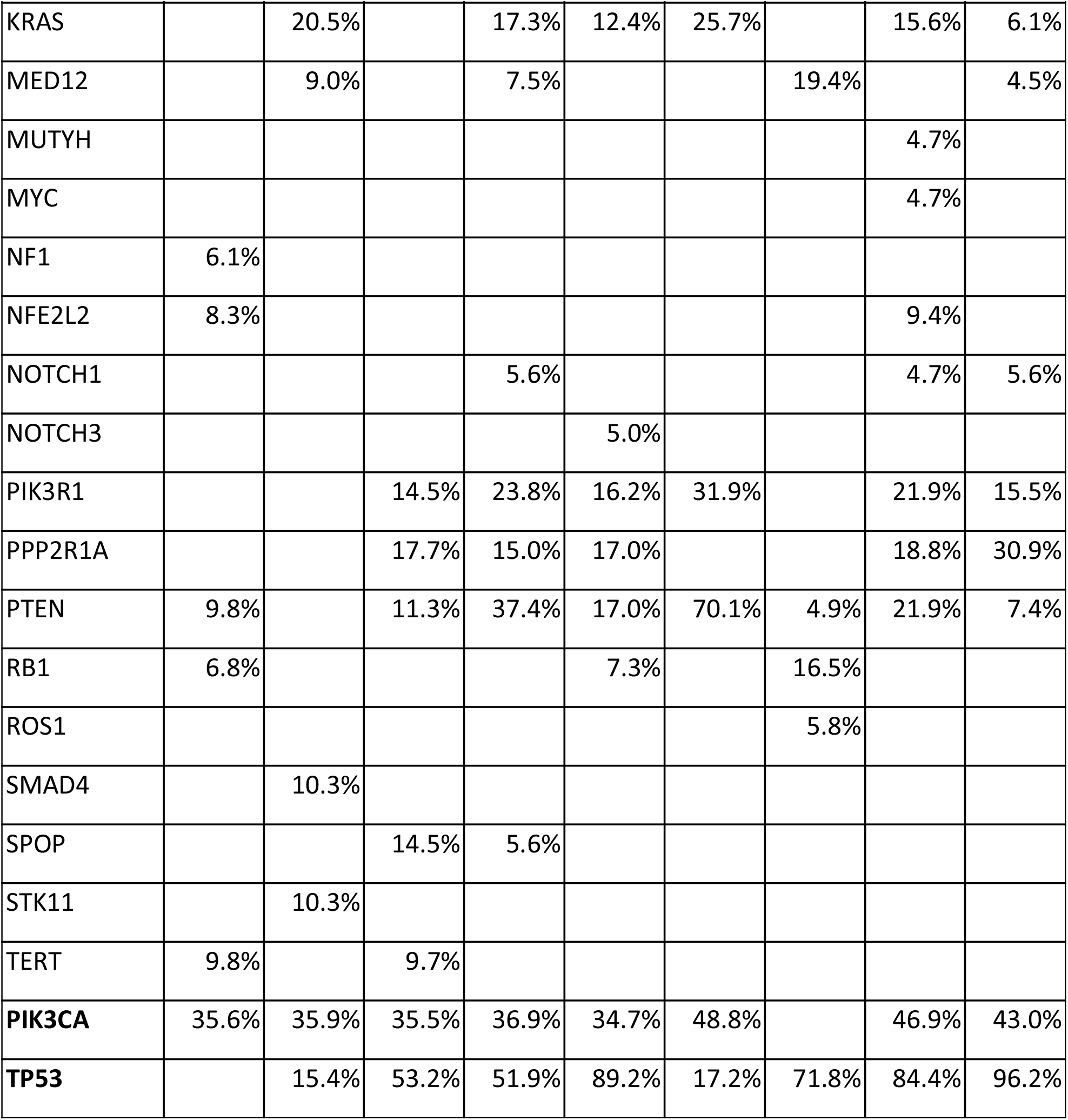
Gene Mutations by Cancer Type Chart

| GENE | CESC | ECAD | UCCC | UCEC | UCS | UEC | ULMS | UMEC | USC |
| --- | --- | --- | --- | --- | --- | --- | --- | --- | --- |
| AKT1 |  |  |  |  |  | 7.9% |  |  |  |
| APC |  | 9.0% |  |  |  |  |  |  |  |
| ARID1A |  | 14.1% | 19.4% | 27.1% | 10.0% | 46.1% | 4.9% | 29.7% | 8.5% |
| ATM |  | 9.0% | 14.5% |  |  | 6.0% |  |  |  |
| ATRX |  |  |  |  |  |  | 38.8% | 4.7% |  |
| BAP1 | 9.1% |  |  |  |  |  |  |  |  |
| BCOR |  |  |  |  |  | 12.5% |  |  |  |
| BRCA2 |  |  |  |  |  |  |  |  | 5.4% |
| CDKN2A |  |  | 8.1% |  |  |  |  |  |  |
| CREBBP |  |  |  |  | 6.2% |  |  |  |  |
| CTCF |  |  |  |  |  | 11.3% |  |  |  |
| CTNNB1 |  |  |  | 16.4% |  | 33.6% |  |  |  |
| DAXX |  |  |  |  |  |  | 5.8% |  |  |
| EP300 | 9.8% |  | 9.7% |  |  |  | 5.8% |  |  |
| ERBB2 |  | 15.4% |  |  |  |  |  |  |  |
| ERBB3 |  | 9.0% | 8.1% |  | 5.8% |  |  | 6.3% | 4.9% |
| ERBB4 |  |  |  |  |  |  | 4.9% |  |  |
| FAT1 | 7.6% | 7.7% |  |  | 5.4% |  | 4.9% |  |  |
| FBXW7 | 13.6% |  | 16.1% | 7.9% | 21.6% | 8.3% |  | 21.9% | 23.1% |
| FGFR2 |  |  |  | 7.9% |  | 11.1% |  |  |  |
| GNAS |  | 9.0% |  |  |  |  |  |  |  |
| HLA-B | 6.8% |  |  |  |  |  |  |  |  |
| KDR |  |  |  |  |  |  | 4.9% |  |  |
| KMT2C | 9.8% |  | 8.1% |  |  |  |  |  | 4.5% |
| KMT2D | 22.0% | 15.4% |  | 8.9% | 10.0% | 7.4% | 6.8% |  | 5.4% |
| KRAS |  | 20.5% |  | 17.3% | 12.4% | 25.7% |  | 15.6% | 6.1% |
| MED12 |  | 9.0% |  | 7.5% |  |  | 19.4% |  | 4.5% |
| MUTYH |  |  |  |  |  |  |  | 4.7% |  |
| MYC |  |  |  |  |  |  |  | 4.7% |  |
| NF1 | 6.1% |  |  |  |  |  |  |  |  |
| NFE2L2 | 8.3% |  |  |  |  |  |  | 9.4% |  |
| NOTCH1 |  |  |  | 5.6% |  |  |  | 4.7% | 5.6% |
| NOTCH3 |  |  |  |  | 5.0% |  |  |  |  |
| PIK3R1 |  |  | 14.5% | 23.8% | 16.2% | 31.9% |  | 21.9% | 15.5% |
| PPP2R1A |  |  | 17.7% | 15.0% | 17.0% |  |  | 18.8% | 30.9% |
| PTEN | 9.8% |  | 11.3% | 37.4% | 17.0% | 70.1% | 4.9% | 21.9% | 7.4% |
| RB1 | 6.8% |  |  |  | 7.3% |  | 16.5% |  |  |
| ROS1 |  |  |  |  |  |  | 5.8% |  |  |
| SMAD4 |  | 10.3% |  |  |  |  |  |  |  |
| SPOP |  |  | 14.5% | 5.6% |  |  |  |  |  |
| STK11 |  | 10.3% |  |  |  |  |  |  |  |
| TERT | 9.8% |  | 9.7% |  |  |  |  |  |  |
| **PIK3CA** | 35.6% | 35.9% | 35.5% | 36.9% | 34.7% | 48.8% |  | 46.9% | 43.0% |
| **TP53** |  | 15.4% | 53.2% | 51.9% | 89.2% | 17.2% | 71.8% | 84.4% | 96.2% |

Tumor suppressor gene TP53 is also mutated in all endometrial cancers at different rates (Table 2) [24] [25][26], but mainly on grade 3 tumors and not on grade 1, indicating that TP53 is implicated on tumor progression but not on tumor initiation as is the case of PTEN.

Type II neoplasms develop even in the absence of unopposed estrogen. These tumors begin with a precursor lesion called Endometrial Intraepithelial Carcinoma (EIC) [27][28]. The most common cancer of this type is Uterine Serous Carcinoma (USC) [29][30], previously named Uterine Papillary Serous Carcinoma [31][32] (Table 1). Patients diagnosed with Type II uterine cancers are usually postmenopausal. The correlation between EIC and USC is the overexpression of mutated p53 protein on both. The gene responsible for the expression of p53 is TP53 (see Table 2). TP53 is a tumor suppressor gene known for being the most frequently mutated gene in all kinds of cancers [33][34]. In our study, only one cancer type, Cervical Squamous Cell Carcinoma (CESC) does not have TP53 in its list of 12 most mutated genes (Table 2). Type II endometrial cancers usually invade the myometrium (see Figure 1). The depth of myometrial invasion, grossly measured as the inner-third, middle-third and outer-third, is associated with metastasis. Different percentages of lymph node and pelvic node metastasis are associated with tumor grade and myometrial invasion depth [35][36].

In Type II uterine cancers, tumor suppressor gene TP53 is mutated in precursor lesions (EIC), which indicates that TP53 is mutated early and thus, is a key driver in the initiation of tumorigenesis.

Neoplasms of Type II are Uterine Clear Cell Carcinoma (UCCC) [37][38], Uterine Carcinosarcoma (UCS) [39][40], Uterine Serous Carcinoma (USC) [41][42], Uterine Mixed Endometrial Carcinoma (UMEC) [43] [44], and others that were not part of this research due to the small number of cases available.

### Cervical Cancers

There are two main cancers of the uterine cervix: Cervical Squamous Cell Carcinoma (CESC) [45] [46], and Endocervical Adenocarcinoma (ECAD) [47][48]. Their mutational profiles are quite different as shown on Table 2. ECAD is the neoplasia on Table 2 with the highest number of unique mutated genes, namely, APC, ERBB2, GNAS, SMAD4, and STK11. The vast majority of malignancies of the cervix are of the squamous cell carcinoma type (96%) and the rest are glandular lesions, or endocervical adenocarcinomas (4%). In most cases (90% or more) these neoplasms begin with a human papillomavirus (HPV) infection [49] [50]. HPV has more than 130 known strains and the particular strains associated with cervical cancers are HPV16 and HPV18 [51][52].

Squamous Cell Carcinoma (CESC) of the uterine cervix starts in a region of the exocervix called the transformation zone (TZ) (See Figure 1). The endocervical canal is lined by two distinctive types of epithelium, squamous and glandular (columnar). The site where the two types of epithelium meet is known as the squamous-columnar junction (SCJ). The SCJ is located at birth in the endocervical canal. This junction moves to the external surface of the cervix facing the vagina after puberty. The zone between the original SCJ and the new SCJ is known as the transformation zone (TZ) where most malignant squamous cell neoplasms develop [53][54]. At the molecular level, some studies show that the most frequently mutated gene is PI3KCA (27.1% of all cases) [55][56] which is in close agreement with out findings (35.6%) as shown on Table 2.

Cervical adenocarcinoma (ECAD) arises and develops in the glandular (columnar) epithelium of the endocervical canal [57]. ECAD in situ, also known as “the usual type” comprises 80% of all adenocarcinoma cases. Other subtypes are: mucinous adenocarcinoma [58], clear cell adenocarcinoma [59], adenosquamous carcinoma [60], and others. These other malignancies were not studied in this research due to the small number of cases reported. As reported by other studies, we found that PI3KCA and KRAS are the most highly mutated genes on ECAD, 35.9% and 20.5% respectively [61][62](Table 2).

## METHODS

Tumor mutational data were obtained from the AACR Project GENIE [1] which has a publicly available set of files that can be downloaded from their website. The full dataset for all cancer types was downloaded and imported into a local SQL Server database for further processing. We explored the data for uterine and cervical cancers and based on the number of cases available, we chose the nine cancers shown on Table 1. The nomenclature used to label the different cancer types was taken from project OncoTree [63].

We first determined the 12 most mutated genes for each cancer type (Table 2) along with the percentage of tumors that show that mutation. Additional filtering was done, looking for tumors with more than 3 mutations on the list of 12 most mutated genes, or at least two mutations, one of which was a unique gene for the corresponding disease. The next step was to find a feature that could be used to train the convolutional neural network (CNN). We selected two features: mutation variant type (see Table 3) and mutation variant classification (see Table 4) [64]. We counted the actual number of variant types and classifications, calculated the relative percentage of the population, and manually assigned a score that was suitable to train the CNN.

**Table 3.** Mutation Variant Types showing the numbers of mutations found in the data sets.

| ID | Variant Type | Description | Count | PCT % | Score |
| --- | --- | --- | --- | --- | --- |
| 1 | SNP | Single nucleotide polymorphism. A substitution in one nucleotide | 231090 | 83.52 | 0.8352 |
| 2 | DEL | Deletion. The removal of nucleotides | 30067 | 10.87 | 0.4444 |
| 3 | INS | Insertion. The addition of nucleotides | 12704 | 4.59 | 0.3555 |
| 4 | DNP | Double nucleotide polymorphism. A substitution in two consecutive nucleotides | 2340 | 0.85 | 0.1111 |
| 5 | ONP | Oligo-nucleotide polymorphism. A substitution in more than three consecutive nucleotides | 486 | 0.18 | 0.1111 |

**Table 4.**
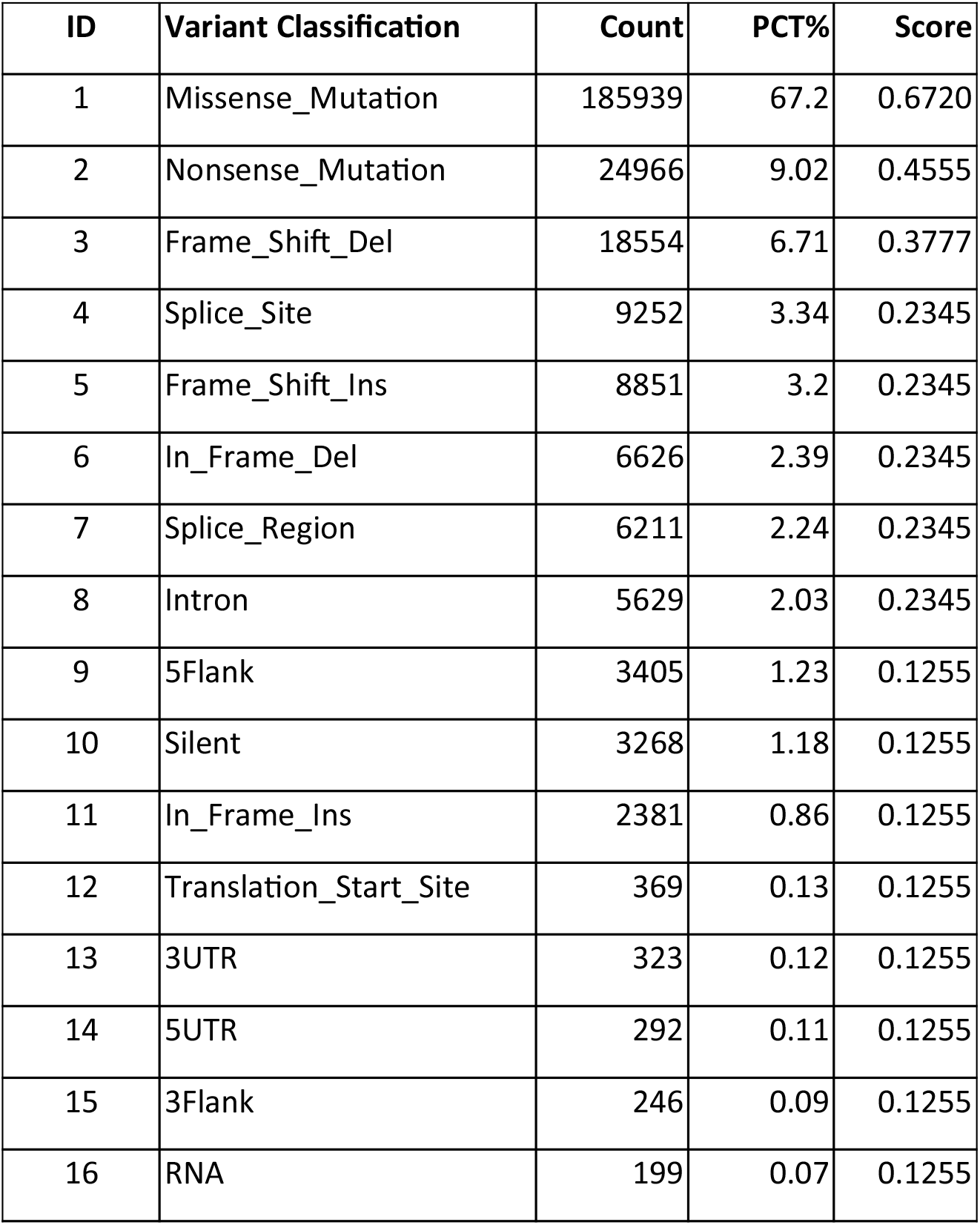
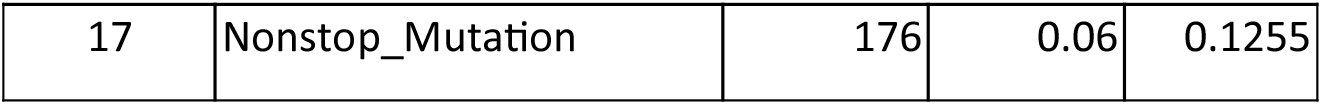
Mutation Classifications

## DEEP LEARNING SOLUTION

Artificial Intelligence (AI), and more specifically, Deep Learning (DL), has been used during the past three decades to solve problems in several areas such as engineering, science, finance, business, social sciences, and others. The solutions are of different kinds: from estimation and prediction to classification, from pattern recognition to natural language processing (NLP). Cancer research is not the exception and several projects have been developed in this area [65][66][67][68].

In this research, a Convolutional Neural Network (CNN) classifier [2] was chosen to classify tumors of gynecological origin. The solution was implemented with a program written in the Python language, making use of the TensorFlow-Keras libraries. The total number of genes was 42 as shown on Table 2, on which, at the bottom, there are 2 genes that were excluded because they are highly mutated in all neoplasias but one, and thus, do not provide any disambiguation information. Those are, oncogene PIK3CA, and tumor suppressor TP53.

For each gene, its variant type and its variant classification scores were used. Since there are 42 genes, 84 data points need to be presented to the CNN input layer. We converted the input 1D vector to a 2D matrix by adding 6 zero-valued dummies at the end. That way, the 90 data points were converted to a 9 by 10 matrix that is fed into the Keras Conv2D input layer. The complete design of the CNN is shown on Figure 2. It has a Conv2D input layer, followed by auxiliary MaxPooling 2D layer, a Dropout layer to remove redundant data, and a 2D to 1D (Flatten) converter layer. The next component is a stack of 4 dense layers, whose neuron numbers and activation functions are shown on Figure 2. The CNN last layer, the output layer, is composed of 9 neurons, each one representing one of the 9 uterine and cervical cancer types. Its activation function is of the type Softmax.

**Figure 2.**
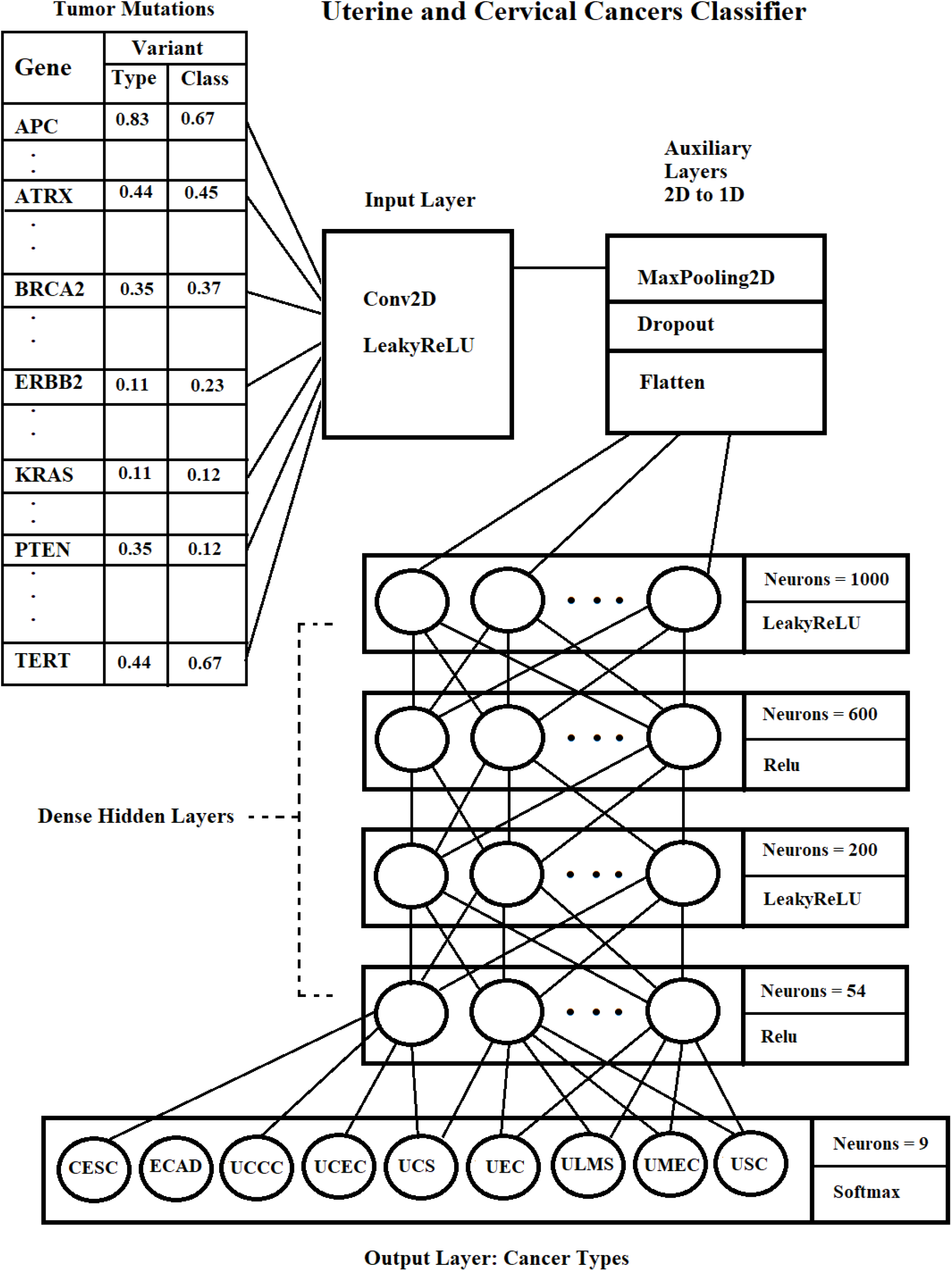
Convolutional Neural Network (CNN)

## RESULTS AND DISCUSSION

The CNN was trained during 120 epochs and in the end, the overall train and test accuracy were 94.3% and 89.2%, respectively. The training accuracy progress during 120 epochs is shown on Figure 3.

**Figure 3.**
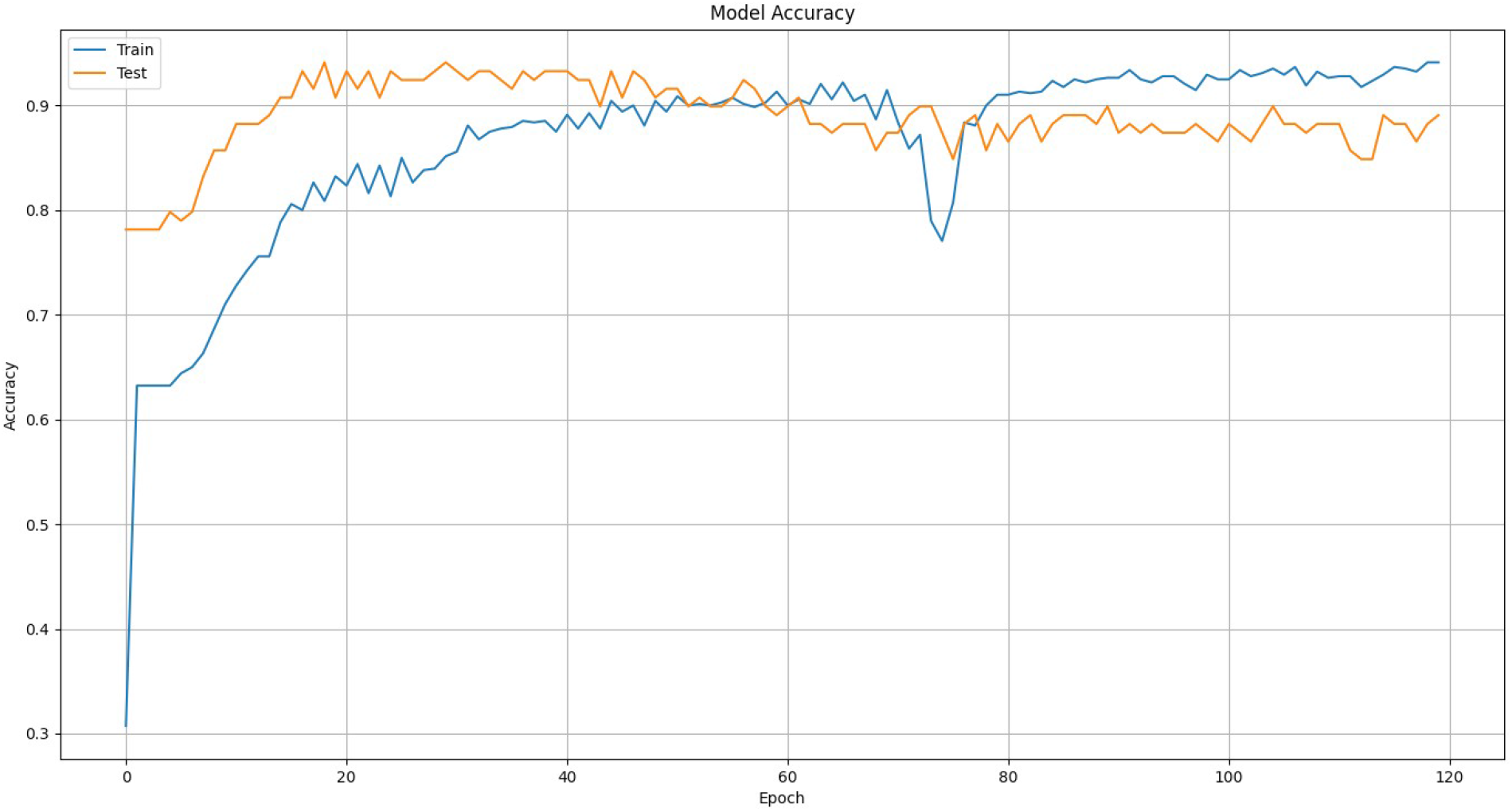
Training Evolution.

Additionally, once the CNN model was saved and ready for evaluation, we run experiments to verify the accuracy of the three datasets, train, test, and evaluation for each of the nine cancer types. The last one, evaluation dataset, was not used during the training process. The number of samples assigned to each dataset, train, test, and evaluation, were 80%, 15%, and 5%, of the whole population, respectively. The results are shown on Table 5. In one case, for cancer type UMEC, the evaluation set consisted of only two tumors that were unsuccessfully classified and thus, the resulting accuracy is zero.

**Table 5.**
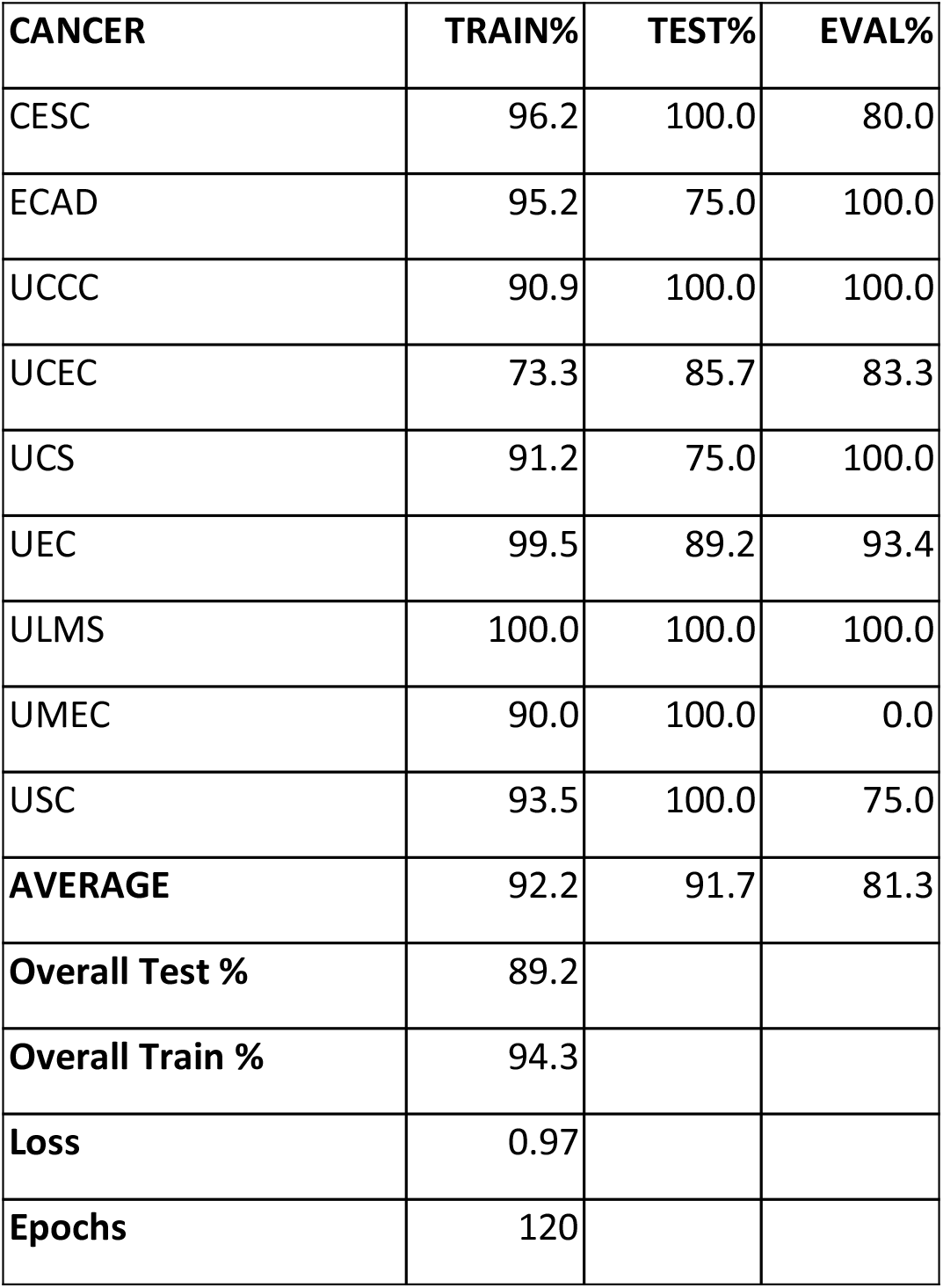
Accuracy by Neoplasia by Data Set

## CONCLUSION

We succeeded in developing a neural network that is capable of accurately classifying tumors of the uterus and uterine cervix based solely on the genetic mutations found on the tumor samples. The resulting overall accuracy is above 90%, which makes this proposed solution a promising tool that should be considered for use in the clinical setting.

